# Genvectors: exploring patterns and unveiling processes driving the geographic distribution of genetic lineages

**DOI:** 10.1101/330761

**Authors:** Leandro Duarte, Gabriel Nakamura, Jacqueline Souza Lima, Renan Maestri, Vanderlei Debastiani, Marcial Quiroga-Carmona, José Alexandre F. Diniz-Filho, Rosane Garcia Collevatti

## Abstract

Sets of local populations show different degrees of gene flow due to dispersal barriers and environmental constraints, which renders genetic composition gradients among populations (genetic turnover). Unveiling biogeographic correlates of genetic turnover is paramount for phylogeography. While some processes (genetic drift, secondary contact) may erase the historical track of genetic turnover, vicariance or ancient dispersal likely leads to genetic divergence among populations. Yet available analyses do not permit direct inference about mechanisms driving genetic turnover. We propose a novel analytical approach called genvector analysis, which fulfills this gap by decomposing genetic compositional dissimilarities between populations based on either haplotypes or other genetic data into genetic eigenvectors. Such procedure allows exploring genetic turnover among sets of local populations, and analyzing their biogeographic correlates based on null model tests. We evaluate the statistical performance of the method on simulated datasets. We also analyzed biogeographic correlates of genetic turnover of *Akodon cursor* in the Brazilian Atlantic Forest. Results revealed that genvector analysis is robust to discriminate biogeographic drivers of genetic turnover. For *Akodon cursor* analysis, we observed that while for the entire species, all predictors considered (except for elevation) explained genetic turnover, within phylogroups some factors varied their importance. Genvector analysis was demonstrated to be useful for several different purposes in phylogeography, and complementary to classic analytical tools widely used by phylogeographers, such as AMOVA or DAPC. The role of ancient versus recent biogeographic events, the relationship between morphological divergence or abiotic variables and genetic turnover can easily be investigated using genvector analysis.

## INTRODUCTION

Local populations show different degrees of connection via gene flow mediated by individual dispersal (Hanski and Gilpin 1991). Given the homogenizing effect of gene flow, the degree of isolation or connectivity among a set of local populations can be properly estimated by evaluating the amount of shared genetic variation among them (Garcia-Verdugo et al. 2010; Murphy et al. 2010). Accordingly, the higher the gene flow between two populations, the lower the dissimilarity between them in terms of changes in genetic composition (hereafter genetic turnover), simply because they will constantly share a greater proportion of genetic variation (Eckert et al. 2008; Templeton 2021). The importance of gene flow for population differentiation can be dynamic, which implies that a local population may be connected to others for a certain time period and later become isolated due to the gradual emergence of biogeographic barriers that impede dispersal until gene flow ceases entirely (Simões et al. 2014). Alternatively, anciently established barriers preventing gene flow may weaken towards the present due to environmental, physical and/or anthropogenic changes in the landscape, facilitating genetic contacts among these formerly isolated populations (Feder et al. 2013). In such cases, while some degree of genetic turnover might still be detected, the signal of genetic divergence is also likely to be reduced (or even erased) as recent admixture between previously disconnected populations increases (Templeton 2021). Other scenarios involve more gradual patterns of genetic differentiation due to geography (i.e., Isolation-by- Distance, IBD) or environment (Isolation-by-Environment, IBE), in which there is no complete cessation of gene flow among populations (Sexton et al. 2014). Rather, a gradual accumulation of local genetic differences is expected due to an inverse association between the rate of interpopulation immigration and geographic distance among populations or environmental dissimilarity among the landscapes in which they occur (Sexton et al. 2014).

Geographic sampling of the genetic composition in sets of local populations informs about the relative levels of genetic turnover among them. Although their genetic composition may be modified by other mechanisms (e.g. locally arisen mutations, non-random mating, selective sweeps), in most circumstances, the genetic turnover observed among populations will primarily depend on the local effects of genetic drift on the distribution of allele frequencies (Sexton et al. 2014; Templeton 2021). Characterizing these compositional changes across geographic space is one of the major goals of phylogeography, a discipline also interested in identifying mechanisms responsible for genetic turnover among local populations (Manel et al. 2003; Storfer et al. 2006; Avise 2009; Balkenhol et al. 2017). These spatial patterns of genetic structuring, which can be expressed as the spatial pattern of distribution of alleles, haplotypes or genotypes, have been analyzed using several approaches in the context of geographical and landscape genetics (see Epperson 2003; Rissler 2016). When these drivers of genetic turnover are proposed as biogeographic factors underlying the genealogical history of populations of a single species, proper hypothesis testing may shed light on populational connectivity from landscape to regional scales (Hand et al. 2016), or even on biogeographic connections across entire regions, biomes or continents (Diniz Filho et al. 2008; Carnaval et al. 2014).

In this context, statistical phylogeographic approaches helped moving that field beyond the essential descriptive nature present in its infancy (Templeton 2004; Papadopoulou and Knowles 2016). For instance, Analysis of Molecular Variance (AMOVA; see Excoffier et al. 1992) has been widely used to analyze dissimilarities in terms of genetic turnover among local populations, based on the proportion of haplotypes shared and those that are exclusively present in only one of them (Fitzpatrick 2009; Maestri et al. 2016; Raffini et al. 2018). AMOVA, which is a permutation procedure akin to approaches often used in ecological studies, such as PERMANOVA (Pillar and Orlóci 1996; Anderson 2001), allows one to evaluate whether pairwise genetic dissimilarities between sets of populations are higher than expected by chance given their within-population dissimilarities. It usually complements exploratory methods to visualize haplotype links and distribution among populations such as haplotype networks (Mardulyn 2012). Nonetheless, AMOVA does not permit direct inference about alternative processes underlying biogeographic correlates of genetic turnover among populations and deeper patterns that have been more explicitly investigated by phylogeographic approaches (e.g., reconstruction of time-calibrated haplotype trees; Templeton 2021). Indeed, despite the ever-increasing number of molecular loci discovered and the development of numerous analytical tools focused on historical biogeography and demographic history of species (Templeton 2004; Epperson 2005; Miller 2005), a more flexible framework capable of simultaneously exploring genetic turnover patterns and inferring the underlying causal mechanisms, while accommodating heterogeneous genetic data structures and spatial predictors, remains a major lacuna in the phylogeographer’s toolkit.

To address these methodological limitations, we introduce here a novel, versatile analytical approach called genvector analysis, based on fuzzy set theory (Zadeh 1965), which allows (1) exploring the distribution of genetic information (e.g. haplotypes) across sets of local populations, and (2) providing a formal statistical framework to test alternative mechanisms driving genetic turnover across geographic space. In classical set theory, the membership function assumes binary values, which means that an element either belongs or does not belong to a given set. In fuzzy set theory, otherwise, any potential element of a set may be described by a membership function defining its degree of belonging to that set ranging between zero and one. Fuzzy sets provide a more realistic categorization of biological entities, such as populations, and have been employed in community ecology and biogeography to define species clusters or composition gradients (Dale 1977; Roberts 1986; Feoli and Zuccarello 1991; Pillar and Orlóci 1991; Olivero et al. 2013). More recently, application of fuzzy set theory has been demonstrated to be useful for the analysis of functional structure of metacommunities (Pillar et al. 2009; Pillar and Duarte 2010), metacommunity phylogenetics (Pillar and Duarte 2010; Duarte et al. 2016; Maestri and Duarte 2020a) and biogeographic regionalization (Maestri and Duarte 2020b). To our knowledge, no attempt to apply fuzzy set theory to explore genetic turnover among single-species populations has been made so far.

We demonstrate the usefulness of genvector analysis for inferring alternative mechanisms driving genetic turnover using simulated datasets, which allowed to build sets of local populations assembled based on known alternative mechanisms (dispersal, secondary contact/genetic drift or vicariance) and then estimate the statistical robustness of genvector analysis for correctly inferring the most likely mechanisms determining genetic turnover and divergence patterns. Finally, we apply genvector analysis on an empirical dataset based on a single-locus marker (haplotypes) to illustrate how the method can be useful for data exploration and biogeographic inference.

## MATERIALS AND METHODS

### Computing pairwise relative genetic covariances to assess genetic turnover among populations

In fuzzy set theory (Zadeh 1965), a fuzzy set denotes the degree of belonging (or membership) of a given element (in the case of this study, an individual), varying between zero and one, to a set. We can easily translate that definition into genetic or evolutionary terms (see also Duarte et al. 2016). Accordingly, the ‘degree of belonging’ can be thought as the amount of relative genetic covariance between a focal haplotype and all other haplotypes in a sample.

Let us start by considering a sample of individuals for which we gathered information on the expression of a single-locus marker, allowing us to address every individual to a haplotype (see Fig. 1 for a numeric example). Noteworthy, this very same logic can be extended to accommodate any genetic data for which a matrix of genetic distances between individuals (matrix **C**, see below) can be calculated. In the so-defined sample, each haplotype shows a variable number of mutational steps from all other haplotypes, which makes it more or less genetically related to every other haplotype in the sample. We start by computing the standardized genetic covariance between two haplotypes *x* and *y* (*c_xy_*), which can be obtained by converting pairwise genetic distances between the haplotypes (e.g. Kimura 1980, 1981) into standardized covariances, as follows:

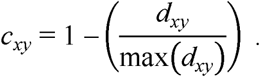

**FIGURE 1.**
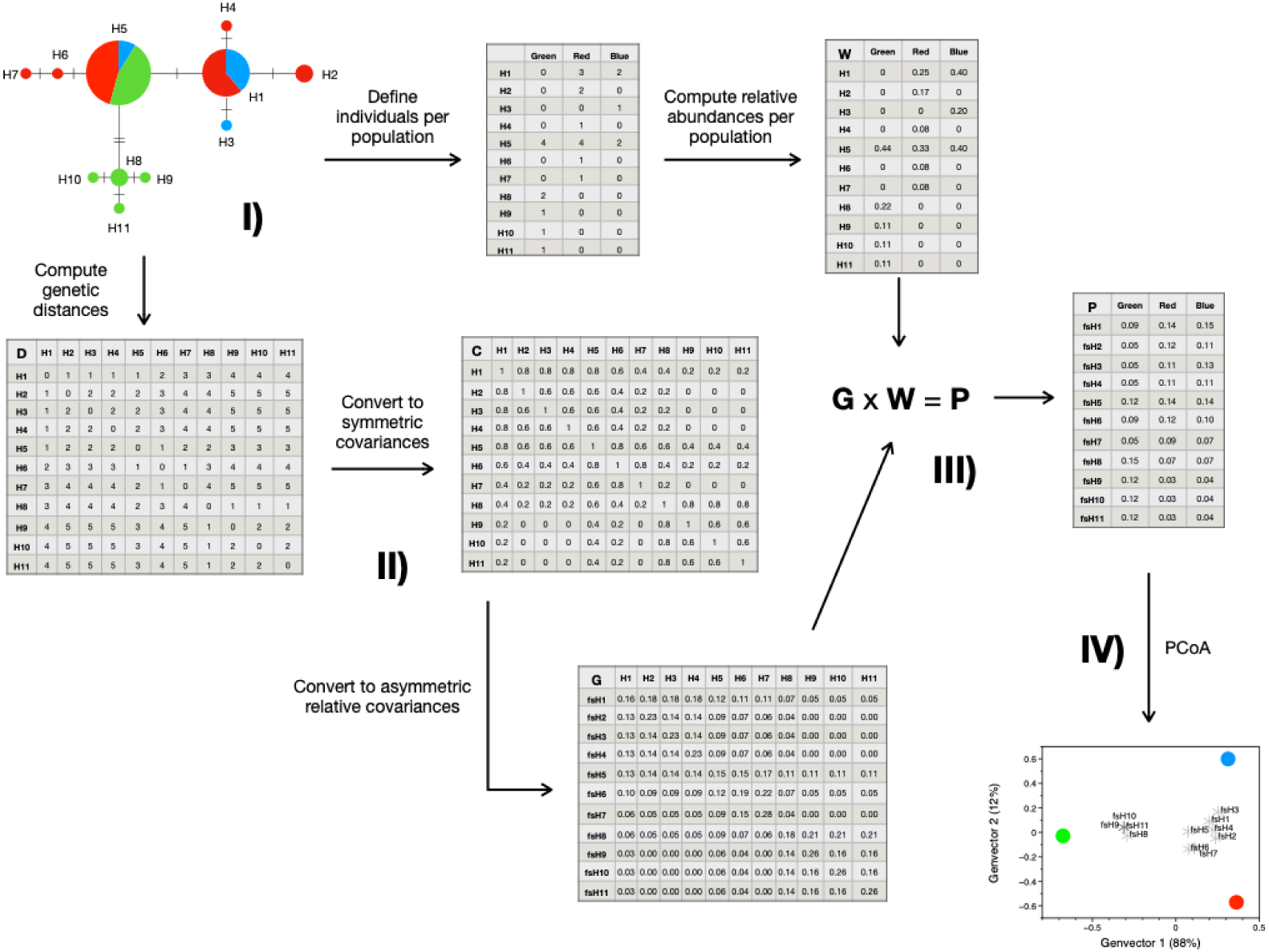
Schematic workflow of genvector computation. I) Network representation of haplotypic patterns built from populations of individuals described by a genetic marker. The network can be easily redescribed in terms of into two input matrices: **D**, genetic distances among haplotypes; computed using an appropriate measure defined by the user (see main text), and **W**, a matrix describing the relative abundance of individuals expressing each haplotype per population. II) The distance matrix **D** is converted to covariance matrix **C**, which is in turn standardized to matrix **G** describing relative genetic covariances between haplotypes. In matrix **G**, the total variance of each haplotype is thereby decomposed into its specific variance and the covariances shared with every other haplotype. III) Matrix **P** is computed by multiplication from matrices **W** and **G**. Matrix **P** describes genetically weighted abundance of haplotypes in each population. IV) Decomposing matrix **P** into eigenvectors by means of principal coordinates analysis generates sets of independent genvectors. Each genvector describes a fraction of genetic variation across the populations. Either dissimilarity matrix **D**_P_ or each independent genvector can be used as response variables to infer biogeographic processes driving genetic turnover.

Accordingly, *d_xy_* is the genetic pairwise distance between haplotypes *x* and *y*, and max (*d_xy_*) is the maximum genetic distance between any pair of haplotypes. Different genetic distance measures implemented in the function ‘dist.dna’ of the package *ape* (Paradis and Schliep 2019) can be used for this purpose. The resulting matrix **C** describes symmetric genetic covariances between haplotypes, which means that *c_xy_* = *c_yx_*. The next step consists in obtaining relative genetic covariances from matrix **C**. By doing so, the covariance *g_xy_* (also known as the ‘degree of belonging’) between any focal haplotype *x* to every haplotype *y* (hereafter defined as a fuzzy set; see Fig. 1) will now depend on the covariance between *y* and all other *z* haplotypes in the sample *(∑c_xz_*, where y z). We can easily compute *g_xy_* as

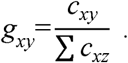

The matrix **G** obtained above describes relative covariances between haplotypes (or fuzzy belongings, see also Pillar and Duarte 2010; Duarte et al. 2016). Matrix **G** will be asymmetric, as it will be defined not only by the pairwise genetic covariance between haplotypes, but also by the topology of the entire haplotype network. Accordingly, ancestral haplotypes will show lower relative covariances to those deriving from it than the opposite (Fig. 1).

Consider now that the abovementioned sample of individuals are distributed across different local populations (represented in Fig. 1 by different colors). As every individual belongs to a corresponding haplotype, we can easily define a matrix describing populations by the proportion of individuals belonging to each haplotype, which we call here matrix **W** (Fig. 1). By multiplication, **G** × **W** = **P**. Matrix **P** describes populations by genetically weighted abundances of haplotypes, which can be used to quantify pairwise genetic turnover between populations by means of widely used resemblance measures, such as Bray-Curtis dissimilarities or Euclidean distances (Legendre and Legendre 2012).

### Computing genvectors

Genetic vectors (or simply genvectors) are the eigenvectors of Principal Coordinates Analysis computed from matrix **P**. The procedure decomposes the total variation in the genetic turnover across the set of populations into independent fractions proportional to its respective eigenvalue λ (Pielou 1984). Each genvector containing a reasonable amount of total variation in **P** (say, 5% or 10%) ordinate populations orthogonally by the amount of genetic resemblance between populations (Duarte 2011). Thus, genvectors enable expressing in few variables the genetic turnover across a set of populations and can be used as an exploratory visualization tool akin to haplotype networks or DAPC plots (Jombart et al. 2010). Moreover, implementing appropriate null models allows using genvector analysis to infer alternative mechanisms driving genetic turnover among local populations distributed across the geographic space. Either exploratory analysis or mechanism inference can be carried out using the function ‘genvectors’ implemented in the R package *Herodotools* (Nakamura et al. 2024).

The function ‘genvectors’ allows alternative input data. If the genetic marker is a single-locus marker (haplotypes), two input datasets are required: (1) a *.fas file containing aligned genetic sequences representing each sampled individual, and (2) a matrix describing the occurrence of every individual (columns) across a set of sites considered as local populations (rows). Based on these two datasets, the function ‘HaploDist’ extracts the haplotypes from the *.fas file using the function ‘haplotype’ implemented in the R package *pegas* (Paradis 2010), and computes the matrix **D** describing all pairwise distances between haplotypes (*d_xy_*, see section *Computing pairwise relative genetic covariances to assess genetic turnover among populations*), using the function ‘dist.dna’ implemented in the R package *ape*, which is then converted to matrix **C** and standardized to obtain matrix **G**. The default option of ‘HaploDist’ computes matrix **D** based on the distance framework derived by Kimura (1980), sometimes referred to as Kimura’s 2-parameters distance (see Paradis and Schliep 2019). Finally, matrix **W** describes local populations by the proportion of individuals belonging to each haplotype, *i.e.* the relative abundance of haplotypes. Matrix **G** (containing relative covariances between haplotypes) is then used to weight the relative abundance of haplotypes per local population (matrix **W**) by their relative genetic covariance, generating matrix **P**, which describes haplotype-based genetic turnover among sets of populations. Submitting matrix **P** to Principal Coordinates Analysis (PCoA) allows computing genvectors. Alternatively, the function ‘genvectors’ also accepts other types of genetic datasets as input, as the user can enter directly as a matrix containing pairwise genetic distances between individuals instead of a *.fas file.

### Inferring biogeographic mechanisms driving genetic turnover

Function ‘genvectors’ allows assessing biogeographic correlates of genetic turnover among sets of populations and inferring the most plausible mechanism driving genetic turnover (unconstrained dispersal, secondary contact/genetic drift, or vicariance). The rationale of mechanism inference in genvector analysis relies on two complementary null models described below:

The first null model is called ‘turnover test’, which was designed to test the hypothesis that the states of a biogeographic factor **E**, which can be categorical or quantitative, explain the genetic turnover among sets of local populations, under the null hypothesis that this factor **E** does not prevent the dispersal of individuals among local populations (Fig. 2). The null model employed to test that hypothesis (*turnover test*) is a classical permutation-based procedure assuming independence between the distribution of haplotypes across local populations and **E**. Pairwise genetic dissimilarities between populations are computed from the matrix containing populations by genetically weighted abundances of haplotypes–**P** (hereafter D**_P_**)–using an appropriate resemblance measure, such as Euclidean distances or Bray-Curtis dissimilarities (Legendre and Legendre 2012). The test consists of 1) computing a F_Obs._ statistic. If D**_P_** is directly modelled on **E**, the test is a dissimilarity regression on distance matrices (hereafter called ADONIS; McArdle and Anderson 2001). For single haplotypic eigenvectors modelled on **E**, the test is an ordinary linear squares (OLS) model; 2) freely permuting the populations a number of times (say 999); 3) at each permutation step, computing F_null_; and 4) defining the probability of obtaining the observed statistic by chance (H_0_ = F_Obs._ ≤ F_null_), as the proportion of permutations in which F_null_ exceeded F_Obs_. Accordingly, if this first hypothesis is not corroborated (*P*_(FObs. ≤ Fnull)_ > 0.05, H_0_ is not rejected), the test allows inferring that the factor **E** does not influence the dispersal of individuals and that they are able to freely disperse across the set of populations without interference, indicating that unconstrained dispersal is the most plausible mechanism determining genetic turnover across space (Fig. 3b). In such cases, no further test is needed.

**FIGURE 2.**
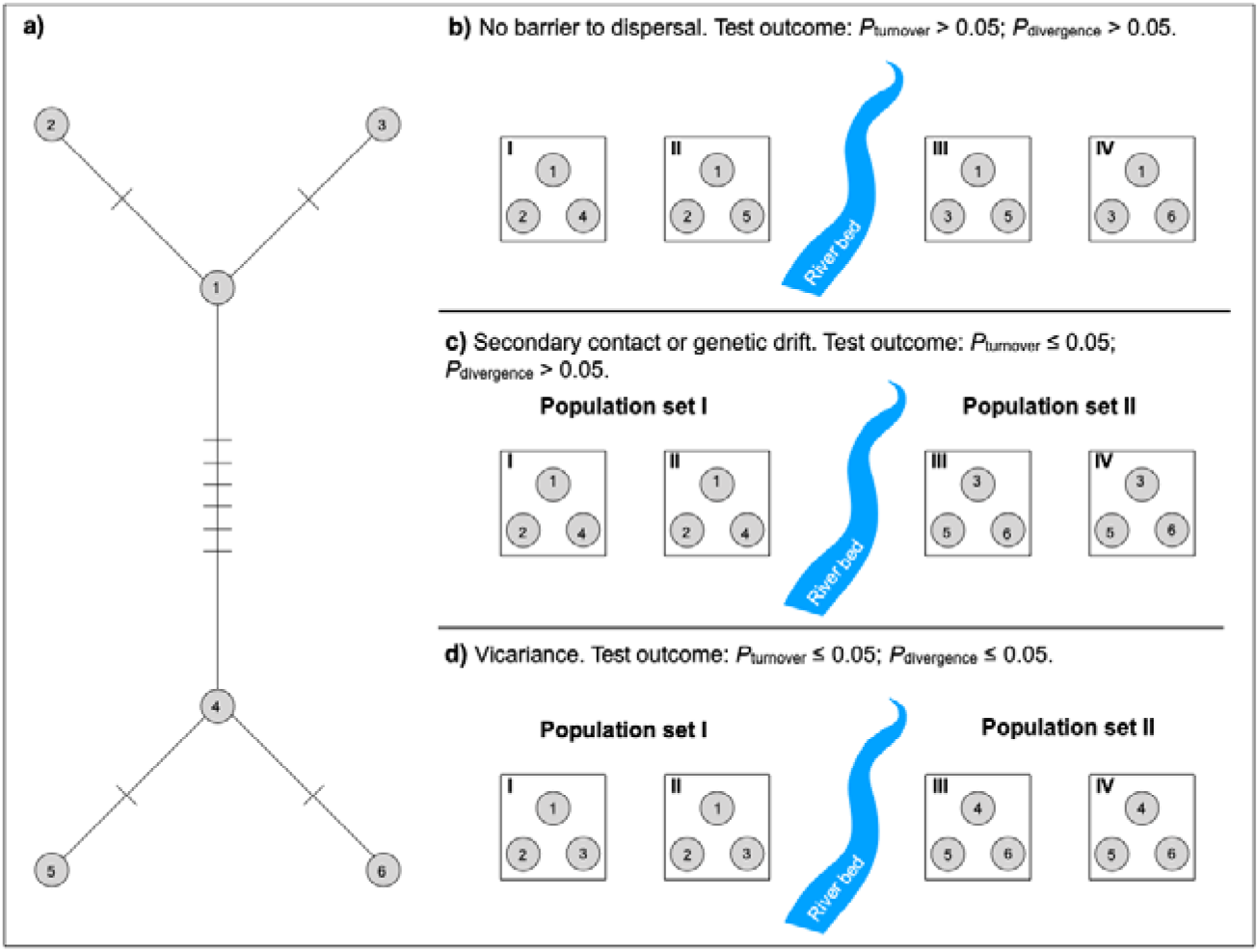
Hypothetical scenarios driving haplotype-based genetic turnover among two sets of populations I-IV located at opposite margins of a river, which may represent a barrier to dispersal. a) Haplotype network showing the number of mutational steps connecting haplotypes 1 to 6. b) Gene flow occurs throughout all populations, which means that the river does not represent a barrier to dispersal among them. Both null model tests implemented in genvector analysis should return *P*-values > 0.05. Therefore, we can conclude that both sets of populations correspond to the same metapopulation. c) Lack of gene flow among sets of populations located at opposite margins of the river. Although haplotype composition is similar between populations located at the same margin of the river (‘turnover’ null model test should return *P*-value _≤_ 0.05), genetic divergence between them is not correlated to the dispersal barrier between them (‘divergence’ null model test is expected to show *P*-value > 0.05). Thus, we can conclude that either gene flow has occurred only recently, as the river is not a barrier to dispersal anymore (secondary contact), or the river has become a barrier to dispersal only recently (genetic drift). d) In this scenario, neither haplotype composition nor genetic divergence between the two sets of populations are correlated to the dispersal barrier between them (the river), indicating lack of gene flow among sets of populations located at opposite margins of the river. Both null model tests should return *P*-values _≤_ 0.05, and we conclude that interruption of gene flow has historical origin, and genetic turnover was most likely caused by vicariance.

**FIGURE 3.**
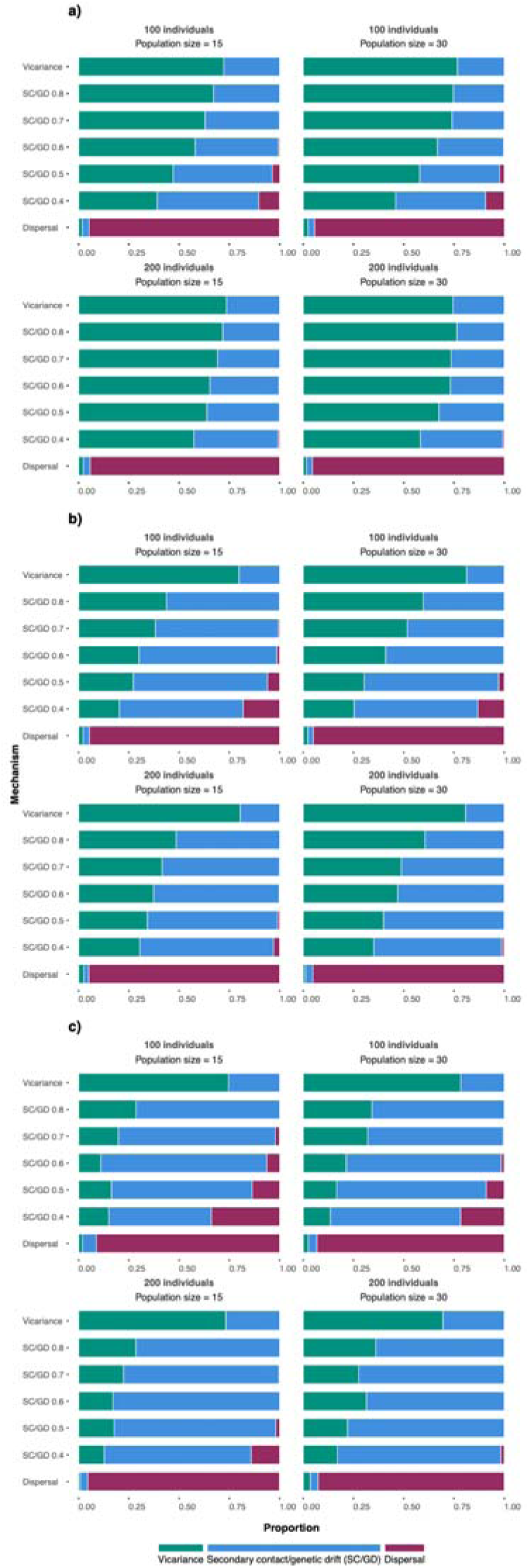
Evaluation of the statistical power of genvector analysis using simulated datasets built using different nominal population sizes, number of individuals per population, and number of states of the biogeographic factor **E** - a) two states, b) three states, and c) four states). For a given dataset simulated based on a predefined mechanism, the corresponding bar indicates the proportion of tests outcomes in terms of process inference. SC/GD 0.4-0.8 refers to the proportion of individuals whose affiliation to a population was preserved during the simulation, ranging between 40% and 80% of the individuals (see main text for details).

Otherwise, whether the null hypothesis of the turnover test is rejected (*P*_(FObs. ≤ Fnull)_ ≤ 0.05), the test allows concluding that populations occurring under similar levels of **E** are more genetically correlated to each other than to populations occurring under different levels of **E** (Fig. 2c, d). Therefore, we may reject unlimited dispersal as the mechanism driving genetic turnover. Rather, gene flow among populations is likely limited or modulated by the barrier represented by the factor **E**. In such scenarios, its influence on genetic turnover can be interpreted based on two alternative mechanisms: secondary contact or genetic drift (Fig. 2c). Considering the first mechanism, either local population originally isolated by the dispersal barrier represented by **E** increase gene flow with each other (secondary contact) and decrease their dissimilarity in terms of genetic composition; considering the second one, local populations formerly connected by dispersal weaken gene flow as a recent dispersal barrier arises and the local effect of genetic drift promotes compositional differences among them. At this point, it is important to recognize that discriminating between these two mechanisms is not trivial, as the turnover pattern generated by either of them is expected to be similar. Therefore, additional analyses, beyond the scope of the method presented here, are necessary to infer between these two mechanisms. We discuss this point in more detail over the next sections; or physical or environmental factors associated with strong vicariance in the past (Fig. 2d). As H_0_ rejection from turnover test itself does not allow discriminating between those two scenarios, carrying out a second null model test is necessary to infer the evolutionary mechanism driving genetic turnover among sets of local populations is needed.

To discriminate between patterns resulting from secondary contact or genetic drift from those caused by vicariance, a second null model called ‘divergence test’ is carried out. This model was designed to test whether the influence of different states of the biogeographic factor **E** on genetic turnover is related to the genetic divergence among them, under the null hypothesis that genetic turnover is independent of the amount of genetic divergence among local populations. Accordingly, a second round of permutations is carried out to compute F_null_. Thus, after computing F_Obs._ (step 1, as described for turnover test), the procedure consists of freely shuffling haplotypes to generate random pairwise genetic covariances among haplotypes (Kembel et al. 2010), while keeping constant the relative abundance of haplotypes in **W**. By doing so, the procedure generates null matrices **C**, **G** and **P**. The procedure is repeated several times (say 999), and, at each permutation procedure, null D**_P_**and, alternatively, null genvectors are computed. In this latter case, null genvectors are submitted to procrustean adjustment (Jackson 1995), and fitted values between observed and null genvectors are obtained. For each permutation step, take null D**_P_**or selected adjusted null genvectors as response variable in ADONIS or OLS (another framework, such as GLS or GLM might be used instead), respectively, using **E** as predictor factor, and compute the corresponding F_null_ values. Finally, generate a set of F_null_ to get the probability under the null hypothesis (H_0_ = F_Obs._ ≤ F_null_). Whenever H_0_ is not rejected (*P*_(FObs. ≤ Fnull)_ > 0.05), we conclude that the association between genetic turnover and **E,** corroborated by the turnover test, is not mediated by the genetic covariances among haplotypes (inferred from the divergence test). In such case, the most plausible mechanisms driving genetic turnover might be either recent secondary contact or genetic drift, which are mechanisms that are either expected to blur the signal of genetic divergence among local populations (Fig. 3c). Otherwise, whether H_0_ is rejected (*P*_(FObs. ≤ Fnull)_ ≤ 0.05), we can conclude that the influence of factor **E** on the genetic turnover across local populations occurring in different regions is proportional to the amount of genetic covariance among them. That is to say that **E** most likely influenced the genetic divergence among sets of populations via longstanding vicariance (Fig. 3d).

### Validating process inference using simulated data

To evaluate the robustness of genvector analysis for mechanism inference, we simulated datasets varying in number of individuals, haplotypes and the number of states in the biogeographic factor **E**. To minimize noises in the simulation process, we included several control checkpoints in the simulation algorithm, which are described over the next sections. Whenever a given simulation failed to pass a checkpoint, it was discarded. To guarantee a minimum number of valid simulations, for each input configuration we carried out 5,000 simulation tries.

### Simulation design

The first step in the simulation process involved defining basic input configuration features: nominal number of individuals (100 or 200), number of populations (15 or 30) and number of states in factor **E** (2, 3 or 4), which was simulated as categories ranging between 0 and 100. For instance, for a simulation where **E** had three states, they were defined as 0, 50 and 100. Those values defined the population’s niche position (*p_j_*). Each local population was addressed to a single state of **E**, indicating that such population was located in a given biogeographic region. Moreover, the assembly process underlying the distribution of individuals across the populations was also previously defined (unlimited dispersal, secondary contact/genetic drift or vicariance).

For each preset input configuration, evolutionary relationships among individuals were simulated as a tree using Grafen’ ρ branch length adjustment = 5 (Grafen 1989; Duarte et al. 2018), which means that evolutionary covariances accelerated towards the tips of the tree. Then, for each tree, individuals were addressed to a given individuals’ pool, which was carried out using a k-means discriminant analysis (Legendre and Legendre 2012) with the number of centers equal to the number of states in **E**. By doing so, we could define which individual evolved in each region defined by **E**. Then we simulated a niche trait (*t_i_*) for every individual evolving according to Ornstein-Uhlenbeck (OU) with multiple optima (θ) model (Hansen et al. 2008; Beaulieu et al. 2012) using the function ‘mvSIM’ of the R package *mvMORPH* (Clavel et al. 2015). Niche optima were set for each group of individuals previously defined, and each one corresponded to a state of **E**. By doing so, we established a link between each geographic region (defined by **E**) and genetic divergence in the individual’s pool.

For simplicity, other parameters of niche trait simulation were kept constant across all simulation configurations (OU’s α = 50; OU’s σ = 10). Having thereby defined the individual’s niche position (*t_i_*) and the site’s niche condition (*p_j_*), we were then able to attribute every individual to a single population by adapting a deterministic Gaussian function proposed by Minchin (1987) and widely used in metacommunity simulation protocols (Peres-Neto et al. 2017; Sokol et al. 2017; Duarte et al. 2023). Accordingly, the affinity of an individual to a given population was computed as follows:

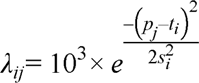

Where *s* is a measure of niche breadth computed for every individual as a Gaussian variable (*N*[10,2]). For every individual, a distribution of affinity values λ_ij_ was computed considering each population. The population for which a given individual showed the maximum λ_ij_ was taken as the population to which the individual belongs. Whenever an individual showed similar affinities for different populations, it was removed from the individual’s pool. In turn, empty populations were also removed from the population pool. Only simulations showing population pools with ≥ 80% of the nominal number of populations were further analyzed. This explains why different sets of valid simulations showed slight differences in the final number of individuals (*x̅*= 97.29 0.43 and = 195.03 8.32 for 100 and 200 nominal number of individuals, respectively) and/or number of populations (*x̅* ± *sd* = 13.55 1.26 and *x̅* ± *sd* = 27.87 1.31 for 15 and 30 nominal number of populations, respectively. See also Table S1). Indeed, in each simulation, only a minor fraction of individuals was discarded from the analyses, as the simulation algorithm often achieved to address the individuals to a single population. These checkpoints guaranteed that every simulation rendered a set of local populations where each individual occurred only in a single population, which, in turn, might contain a variable number of individuals.

After defining populations of individuals across the regions, we simulated genetic sequences evolving under the HKY model of evolution and containing 1000 nucleotides for each individual using the function ‘seqgen’ (Rambaut and Grass 1997) of R package *phyclust* (Chen 2010). Those sequences where further converted into a *.fas file, which was then used as input in ‘genvectors’ function. Note that closely related individuals shared similar niche positions (*t_i_*). As the attribution of individuals to populations was determined by the match between individual’s niche position and site’s niche condition characteristic of given region, we guaranteed that biogeographic regions in **E** represented true dispersal barriers determining genetic turnover.

### Evaluating statistical properties of genvector analysis

As explained above, data simulation was designed to express strong association between a biogeographic factor **E** and genetic resemblance among local populations occurring in a same region (as we should expect to occur when IBD, IBE, or vicariance are the mechanisms generating genetic structure). To get a scenario where **E** was not associated with limited gene flow across local populations (unlimited dispersal scenario), we simply randomly shuffled the original affiliation of each population to a state in factor **E** prior to genvector analysis. Such **E**_0_ factor was then taken as the biogeographic factor in the analysis, while the original simulated matrix **P** kept unchanged. To get a scenario where only part of the individuals dispersed across the geographic space, somehow overcoming the dispersal limitation imposed by **E** (secondary contact/genetic drift scenario), we shuffled a varying fraction of individuals in a way that 40%, 50%, 60%, 70% or 80% of individuals remain in their native population while the remaining dispersed across the space prior to computing matrix **P**. Genvector analysis was therefore carried out using **E** as predictor and the partially shuffled matrix **P** as response variable. Finally, to get the vicariance scenario, we kept the original simulated datasets (**P** and **E**) to perform genvector analysis, as the simulation design assumed vicariance as the underlying assembly process.

After carrying out 5,000 simulation tries for each combination of assembly scenario (vicariance, genetic drift/secondary contact), number of individuals (100 or 200), number of populations (15 or 30) and number of states in **E** (one, two or three), we performed genvector analysis only for valid simulation tries. For those tries we computed the frequency of outcomes correctly inferring the mechanism determining genetic turnover among local populations, which was taken as an estimate of the statistical power (1 – type II error) of the analysis for detecting the mechanism driving genetic compositional turnover across sets of local populations. As our analytical framework does not have a true null hypothesis, but three alternative ones (dispersal, genetic drift/ secondary contact and vicariance), evaluating statistical type I error does not make sense in our analytical framework (Sokal and Rohlf 1994).

### Applying genvector analysis to an empirical dataset

We demonstrate the applications of ‘genvectors’ in phylogeographic assessments using an empirical dataset of 389 sequences of the cytochrome b (cyt-b) gene that were previously analyzed by Cruz et al. (2025) to address the phylogeographic structure and intraspecific diversification in the Neotropical sigmodontine rodent *Akodon cursor*. Sequences were obtained from individuals captured at 90 localities that widely covered the geographic range of the species across the Brazilian Atlantic Forest, also including populations from ecotonal areas with the Caatinga and Cerrado ecosystems (Cruz et al. 2025). Sequences consisted of fragments of 1,140 bp that represent the entire mitochondrial cyt-b gene. These were retrieved from GenBank using their accession codes and the package *SeqinR* (Charif and Lobry 2007). Considering that previous studies have emphasized the importance of river systems as a main factor driving the genetic differentiation of populations of *A*. *cursor*, we used genvectors analyses to assess whether the spatial distribution of cyt-b haplotypes might be related to the geographic areas delimited by river basins at two hierarchical spatial scales, macro and meso river basins (IBGE 2021). In addition, the role of geographic distances among local populations in shaping genetic differences among localities (i.e., isolation by distance) was also assessed by performing analyses in which latitude and longitude were included as interactive and continuous predictors of genetic turnover. Finally, the potential role of elevation in shaping the species’ phylogeographic structure was evaluated, given that *A*. *cursor* thrives across a wide altitudinal gradient that spreads from sea level to 1,170 m (Cruz et al. 2025).

The analyses were conducted at the species level using the entire set of aligned cyt-b sequences but also splitting this dataset into separate subsets corresponding to the intraspecific phylogroups identified by Cruz et al. (2025) to evaluate the importance of same factors in structuring phylogeographic patterns at smaller scales. Thus, tests were also conducted using a dataset built integrating the haplotypes and localities of the central and Southern phylogroups, and using each one separately. These two groups were selected because they have the highest number of sampled localities, sampled sequences, and showed higher levels of reciprocal geographic overlap. The Northern clade was not analyzed accordingly because it had the fewest sequences and sampled localities. For ADONIS, we computed matrix **P** using frequencies of haplotypes per site (matrix **W**) and square-rooted Hamming distances between haplotypes (matrix **D**). Haplotypic dissimilarities between sites were computed using Bray-Curtis dissimilarities. For OLS, we first computed haplotype-based genvectors based on D_P_ and the first two genvectors, which individually contain more than 5% of total information in **P** (see Table 1), were taken as response variables in two linear models, while the corresponding **E** was used as predictor.

**TABLE 1.** Results of ADONIS- and OLS-based null model tests implemented in genvector analysis for mitochondrial phylogroups of *Akodon cursor* in the Brazilian Atlantic. Tests were conducted to assess the influence of geographic partitioning defined by hydrographic basins, as well as the effects of geographic distance (according to e and longitude coordinates) among populations and altitudinal distribution on the phylogeographic structure observed in sampled populations (Figure S1). For each is, the observed F-value (F_Obs_) and the *P*-values corresponding to the turnover tests (*P*_tur_) and divergence tests (*P*_div_) are provided. OLS-based analyses were performed the first two genvectors (GV), which were computed based on Dp (pairwise haplotype dissimilarities among sets of populations computed from matrix **P**).

| Factors | Models |  |
| --- | --- | --- |
|  | ADONIS | OLS |
| *Entire cyt-b dataset* |  |  |
| Meso Basins | $F_{\text{Obs.}} = 15.57, P_{\text{tur}} = 0.001, P_{\text{div}} = 0.001$ | GV1(56%): $F_{\text{Obs.}} = 18.02, P_{\text{tur}} = 0.01, P_{\text{div}} = 0.03$; GV2(16%): $F_{\text{Obs.}} = 42.23, P_{\text{tur}} = 0.01, P_{\text{div}} = 0.01$ |
| Macro Basins | $F_{\text{Obs.}} = 42.60, P_{\text{tur}} = 0.001, P_{\text{div}} = 0.001$ | GV1(56%): $F_{\text{Obs.}} = 116.92, P_{\text{tur}} = 0.001, P_{\text{div}} = 0.001$; GV2(16%): $F_{\text{Obs.}} = 3.15, P_{\text{tur}} = 0.071, P_{\text{div}} = 0.259$ |
| Latitude*Longitude | $F_{\text{Obs.}} = 5.25, P_{\text{tur}} = 0.002, P_{\text{div}} = 0.009$ | GV1(56%): $F_{\text{Obs.}} = 87.03, P_{\text{tur}} = 0.001, P_{\text{div}} = 0.001$; GV2(16%): $F_{\text{Obs.}} = 4.26, P_{\text{tur}} = 0.011, P_{\text{div}} = 0.056$ |
| Elevation | $F_{\text{Obs.}} = 2.06, P_{\text{tur}} = 0.099, P_{\text{div}} = 0.121$ | GV1(56%): $F_{\text{Obs.}} = 2.56, P_{\text{tur}} = 0.105, P_{\text{div}} = 0.156$; GV2(16%): $F_{\text{Obs.}} = 0.82, P_{\text{tur}} = 0.365, P_{\text{div}} = 0.485$ |
| *Central+Southern phylogroups* |  |  |
| Meso Basins | $F_{\text{Obs.}} = 33.38, P_{\text{tur}} = 0.001, P_{\text{div}} = 0.001$ | GV1(57%): $F_{\text{Obs.}} = 87.57, P_{\text{tur}} = 0.01, P_{\text{div}} = 0.03$; GV2(8%): $F_{\text{Obs.}} = 1.30, P_{\text{tur}} = 0.262, P_{\text{div}} = 0.318$ |
| Macro Basins | $F_{\text{Obs.}} = 7.11, P_{\text{tur}} = 0.001, P_{\text{div}} = 0.005$ | GV1(57%): $F_{\text{Obs.}} = 12.76, P_{\text{tur}} = 0.001, P_{\text{div}} = 0.007$; GV2(8%): $F_{\text{Obs.}} = 0.71, P_{\text{tur}} = 0.393, P_{\text{div}} = 0.443$ |
| Latitude*Longitude | $F_{\text{Obs.}} = 1.25, P_{\text{tur}} = 0.247, P_{\text{div}} = 0.473$ | GV1(57%): $F_{\text{Obs.}} = 10.84, P_{\text{tur}} = 0.002, P_{\text{div}} = 0.006$; GV2(8%): $F_{\text{Obs.}} = 1.15, P_{\text{tur}} = 0.362, P_{\text{div}} = 0.495$ |
| Elevation | $F_{\text{Obs.}} = 0.77, P_{\text{tur}} = 0.473, P_{\text{div}} = 0.737$ | GV1(57%): $F_{\text{Obs.}} = 0.05, P_{\text{tur}} = 0.814, P_{\text{div}} = 0.843$; GV2(8%): $F_{\text{Obs.}} = 0.32, P_{\text{tur}} = 0.583, P_{\text{div}} = 0.786$ |
| *Central phylogroup* |  |  |
| Meso Basins | $F_{\text{Obs.}} = 5.76, P_{\text{tur}} = 0.001, P_{\text{div}} = 0.001$ | GV1(28%): $F_{\text{Obs.}} = 32.69, P_{\text{tur}} = 0.001, P_{\text{div}} = 0.001$; GV2(26%): $F_{\text{Obs.}} = 1.49, P_{\text{tur}} = 0.244, P_{\text{div}} = 0.299$ |
| Macro Basins | $F_{\text{Obs.}} = 1.16, P_{\text{tur}} = 0.291, P_{\text{div}} = 0.586$ | GV1(28%): $F_{\text{Obs.}} = 3.18, P_{\text{tur}} = 0.084, P_{\text{div}} = 0.179$; GV2(26%): $F_{\text{Obs.}} = 0.01, P_{\text{tur}} = 0.908, P_{\text{div}} = 0.951$ |
| Latitude*Longitude | $F_{\text{Obs.}} = 4.51, P_{\text{tur}} = 0.002, P_{\text{div}} = 0.007$ | GV1(28%): $F_{\text{Obs.}} = 10.48, P_{\text{tur}} = 0.001, P_{\text{div}} = 0.007$; GV2(26%): $F_{\text{Obs.}} = 27.68, P_{\text{tur}} = 0.001, P_{\text{div}} = 0.003$ |
| Elevation | $F_{\text{Obs.}} = 1.99, P_{\text{tur}} = 0.049, P_{\text{div}} = 0.039$ | GV1(28%): $F_{\text{Obs.}} = 0.97, P_{\text{tur}} = 0.309, P_{\text{div}} = 0.373$; GV2(26%): $F_{\text{Obs.}} = 6.65, P_{\text{tur}} = 0.011, P_{\text{div}} = 0.022$ |
| *Southern phylogroup* |  |  |
| Meso Basins | $F_{\text{Obs.}} = 2.89, P_{\text{tur}} = 0.009, P_{\text{div}} = 0.012$ | GV1(30%): $F_{\text{Obs.}} = 4.69, P_{\text{tur}} = 0.029, P_{\text{div}} = 0.049$; GV2(12%): $F_{\text{Obs.}} = 5.31, P_{\text{tur}} = 0.038, P_{\text{div}} = 0.044$ |
| Macro Basins | $F_{\text{Obs.}} = 1.47, P_{\text{tur}} = 0.088, P_{\text{div}} = 0.168$ | GV1(30%): $F_{\text{Obs.}} = 2.67, P_{\text{tur}} = 0.101, P_{\text{div}} = 0.128$; GV2(12%): $F_{\text{Obs.}} = 0.18, P_{\text{tur}} = 0.626, P_{\text{div}} = 0.522$ |
| Latitude*Longitude | $F_{\text{Obs.}} = 2.18, P_{\text{tur}} = 0.027, P_{\text{div}} = 0.034$ | GV1(30%): $F_{\text{Obs.}} = 3.80, P_{\text{tur}} = 0.013, P_{\text{div}} = 0.055$; GV2(12%): $F_{\text{Obs.}} = 6.04, P_{\text{tur}} = 0.009, P_{\text{div}} = 0.011$ |
| Elevation | $F_{\text{Obs.}} = 1.63, P_{\text{tur}} = 0.099, P_{\text{div}} = 0.154$ | GV1(30%): $F_{\text{Obs.}} = 0.18, P_{\text{tur}} = 0.663, P_{\text{div}} = 0.722$; GV2(12%): $F_{\text{Obs.}} = 7.19, P_{\text{tur}} = 0.011, P_{\text{div}} = 0.017$ |

## RESULTS

### Simulated dataset

The number of valid simulations out of 5,000 tries varied mostly in relation to the number of states of **E** (Table S2). For the two states scenario, we got, on average, 2,335 valid simulations (*sd* = 132.7) for each input configuration, while for the three and four states scenarios these numbers decreased to 875 (*sd* = 73) and 202 (*sd* = 37.4), respectively.

Overall results for simulation-based analyses are summarized in Figure 3. In general, we found that the number of states of the biogeographic factor affected the robustness of genvector analysis for inferring biogeographic processes (Fig. 3, Table S2). While for dispersal and vicariance the analysis showed to be robust for all simulation scenarios, with statistical power ranging between 0.93 and 0.95 for the dispersal scenario and between 0.74 and 0.81 for vicariance, the same was not observed for the secondary contact/genetic drift scenario. In the two states scenario, secondary contact/genetic drift simulations showed very low statistical power (0.28-0.45). In such simulations, genvector analysis often found vicariance as the biogeographic factor determining genetic turnover among populations. In the three states scenario, statistical power increased in simulations where a larger fraction of individuals was allowed to disperse across the biogeographic barrier (Fig. 3, Table S2). Finally, in the four states scenario, the secondary contact scenario/genetic drift showed appropriate statistical power (ranging between 0.68 and 0.78).

### Empirical dataset

The use of the ADONIS model showed that the scale at which analyses are conducted is important for understanding the turnover and divergence of mitochondrial variation sampled within *Akodon cursor*. While, for the entire species, all predictors considered (except elevation) explained both genetic turnover and divergence, at smaller scales (i.e., within phylogroups) these factors may either loose or gain importance (Table 1). This is evident in the results obtained when analyzing the dataset integrating populations from the central and Southern phylogroups, where latitude and longitude were not relevant (*P*_turnover_ = 0.247, *P*_divergence_ = 0.473). A similar pattern was observed in the evaluations of macro river basin when populations of the central and Southern phylogroups were analyzed separately. In these analyses, elevation was relevant for structuring haplotypic variation of the central phylogroup (*P*_turnover_ = 0.049, *P*_divergence_ = 0.039). Regarding the use of OLS models, their implementation showed similar results concerning the relevance of the scale at which analyses are conducted. Those performed on the complete species were generally consistent with the ADONIS results, with differences arising only when the second genvector was analyzed, as the macro river basin predictor was not associated with this portion of haplotypic variation (Table 1, Fig. 4). A similar pattern was observed in analyses conducted on the dataset integrating the central and Southern phylogroups, where the second genvector was not associated with any predictor factor (*P*_Fobs_ ≥ 0.05). In analyses performed separately on each phylogroup, the lack of association with macro river basins was recurrently obtained and elevation showed association with the second genvector.

**FIGURE 4.**
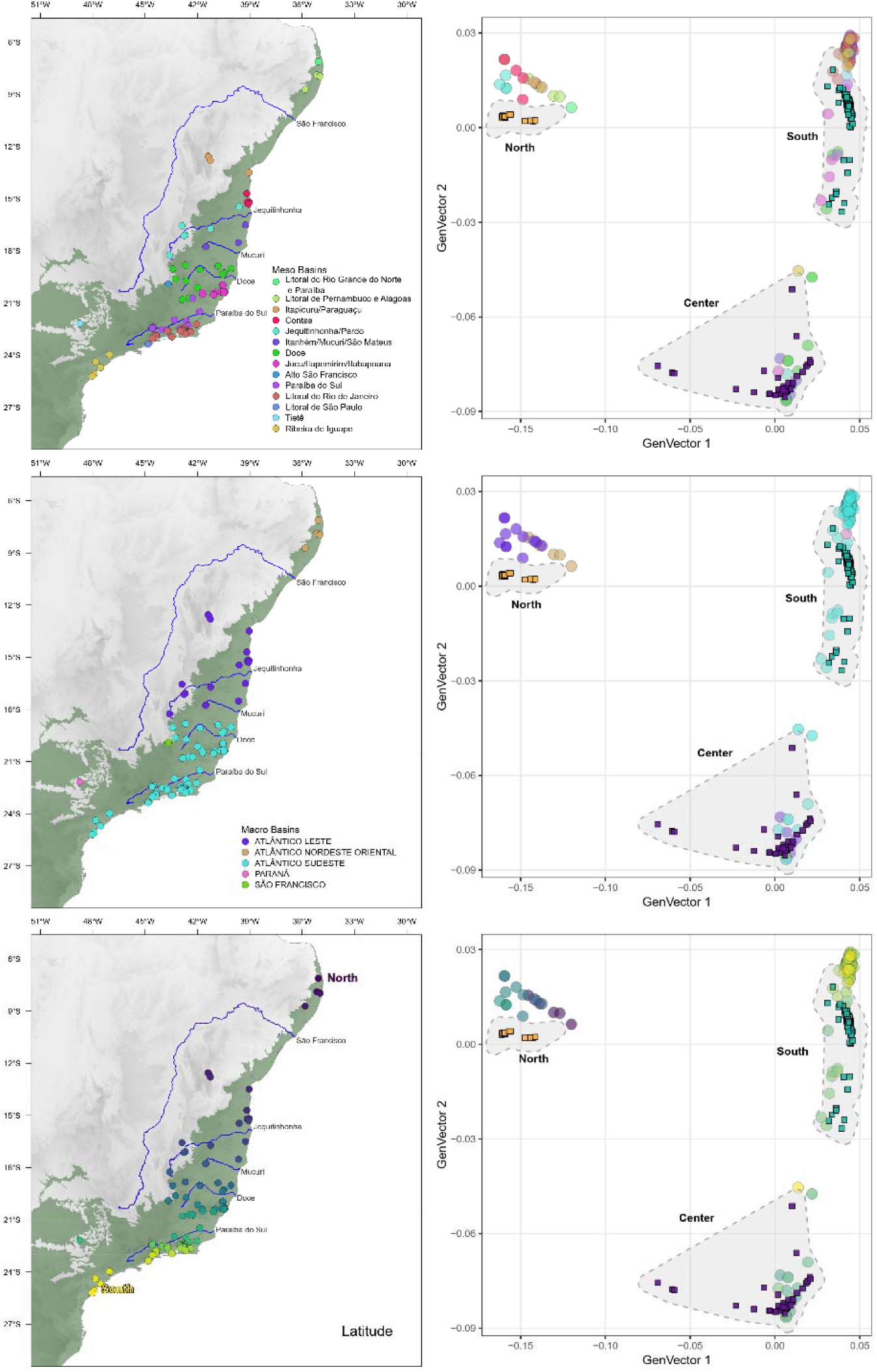
Scatterplots of haplotypes projected onto the first two genvectors calculated in the analyses performed for *Akodon cursor*. Left panels show the geographic distribution of populations and their ordination according to the factors associated with haplotypic structure of populations (see Table 1). Right panels show the distribution of the 90 localities (circles) where individuals of *Akodon cursor* were captured, overlaid with the distribution of the three haplotype groups (or phylogroups, colored squares) identified by Cruz et al. (2025). These phylogroups were also identified as strongly supported clades through phylogenetic analyses based on cyt-b sequences (see Material and Methods).

## DISCUSSION

In disciplines such as phylogeography and population genetics, statistical methods have been implemented to explore and describe patterns of genetic structure among natural populations. Approaches such as Analysis of Molecular Variance (AMOVA; Excoffier et al. 1992) and Discriminant Analysis of Principal Components (DAPC; Jombart et al. 2010) have been widely used for this purpose and to attempt to associate the observed patterns of genetic structure with features of the landscape across which populations are distributed. Indeed, exploring patterns of genetic turnover among sets of populations based on genvectors also reveals clustering patterns among populations, in a similar way to those previously uncovered by classical methods (Hand et al. 2016). However, unlike AMOVA and DAPC, genvector analysis explicitly allows for the integration of spatially structured landscape factors (e.g., elevation, precipitation, temperature) or biogeographic scenarios (e.g., river barriers) as potential drivers of genetic turnover among local populations, and here lies the main advantage of genvector analysis over other available methods. At the same time, it is also worth considering that, despite the statistical ability of genvector analyses to correctly detect patterns of genetic structuring resulting from vicariance scenarios, their main limitation arises when the mechanism responsible for genetic turnover among populations is secondary contact or the local effect of genetic drift. Under these circumstances, it may be useful to contrast results from genvectors analysis with those obtained from other approaches, such as reconstructions of the historical demography of populations and their phylogenetic relationships, which may provide a temporal and directional dimension to the patterns of genetic structure already investigated (Avise 2009; Templeton 2021). Even simple aspects such as considering nucleotide or haplotypic diversity values may help improve the interpretation of these results.

The empirical dataset obtained from the study performed by Cruz et al. (2025) which was used to test the performance of genvector analyses, had previously been evaluated with several analyses based on AMOVA approach to explore the role of the Jequitinhonha River as biogeographic barrier and the existence of two or three genetic clusters within *Akodon cursor*. In all of these implementations of AMOVA analyses, the categorization used as an independent variable does not explicitly account for spatially structured environmental or physical factors possibly driving the distribution of local populations; rather, as is usual when this approach is used (Garcia-Verdugo et al. 2010; Murphy et al. 2010), individuals of each these population were assigned to the groups whose reliability is being evaluated based on the relative amount of genetic variation accumulated by each group. Although these analyses did confirm patterns of genetic turnover consistent with those hypothesized in each proposed scenario (e.g., latitudinal structuration), by themselves they cannot identify the direct effects of more nuanced factors (see Table 1) operating at different scales on genetic turnover to induce patterns of genetic structuration. One aspect that could be effectively addressed through analyses based on the genvector approach, which allow a clear assessment of the influence modulated by different scales, particularly when examining values of F_Obs_ (Table 1).

When genvector analyses were performed using the to the total mitochondrial variation sampled in *Akodon cursor*, it becomes evident that genetic turnover, although associated with both scales of river basins, is more strongly influenced by the scale of macro basins, whereas this influence decreases at smaller scales. This suggests that the genetic turnover of *A. cursor* is spatially structured on a scale that corresponds more closely to the spatial dimensions of macro basins, whereas this pattern of structure is not maintained at smaller spatial scales. The greater relevance of meso basins becomes evident when analyses focus on the genetic variation contained by each phylogroup, either analyzed separately or jointly (see Table 1). A similar pattern is observed when examining the combined effects of geographic coordinates (latitude and longitude) and elevation. In these cases, when analyzing the central and northern groups jointly, geographic distance does not structure haplotypic variation, in contrast to the full species dataset or when the central and southern groups are analyzed separately. Elevation only represented a relevant factor for the genetic structure of the central group, which could be expected considering that the region where this phylogroup is distributed concentrates the greatest variability in this factor (Figure S1).

Finally, it is also important to point out that, although we have demonstrated the usefulness of genvector analysis to investigate the association between genetic turnover and landscape factors in populations spatially dispersed across a multifactorial environment, focusing on neutral loci, the analytical procedures described here may be easily applied to investigate patterns of structure among loci under selection, which may open totally new avenues for future research in fields such as landscape genomics (Kirk and Freeland 2011; Schoville et al. 2012; Edwards et al. 2022). For instance, null model tests implemented in genvector analysis might be used explore associations between non-neutral genetic data and climate change to investigate the influence of environmental fluctuations on the spatial distribution of adaptive genetic variability across sets of local populations (Schoville et al. 2012; Sork et al. 2013). This would allow new insights into how biogeographic settings and landscape dimensions, as well as different geographic scales, have driven and continue to drive selection-based genetic turnover across sets of populations.

Genvector analysis allows to explore genetic turnover across sets of populations and infer the role of alternative mechanisms (unlimited dispersal, secondary contact/genetic drift or vicariance on phylogeographic patterns by means of appropriate null model tests. Moreover, genvector analysis was demonstrated to be a flexible tool in the sense it accommodates to different genetic data types, given these data express evolutionary relationships. These features make the tool useful for several different purposes in phylogeography, and complementary to classic analytical tools widely used by phylogeographers, such as AMOVA (Excoffier et al. 1992), individual-based Bayesian clustering (Pritchard et al. 2000), or DAPC (Jombart et al. 2010). Fundamental questions, such as the role of ancient versus recent biogeographic events, the relationship between morphological divergence and genetic turnover and divergence, and the effects of abiotic variables on genetic variation can easily be investigated using genvector analysis.

## ACKNOWLEDGEMENTS

The authors thank Jorge Sebastião Bernardo-Silva for valuable discussion on the topic addressed in this paper, and Maria Gabriela Junqueira for the help with plot coding. LD, RM, JAFDF and RGC have continuously been supported by CNPq productivity grants, which we gratefully acknowledge. JSL and MQC were supported by CNPq fellowships (grants 439051/2016-9 and 381499/2026-0). LD, GN, RM, VD, MQC and JAFDF are members of the National Institute for Science and Technology (INCT) in Ecology, Evolution and Biodiversity Conservation, supported by MCTIC/CNPq (proc. 409197/2024-6) and FAPEG (proc. 201810267000023).

The authors have no conflict of interests to declare.

## DATA ACCESSIBILITY AND BENEFIT-SHARING

The empirical data used in the analysis are openly available in GenBank database and can be downloaded using corresponding accessing codes available at https://zenodo.org/records/20983800.

## AUTHOR CONTRIBUTIONS

LD, JSL, RGC designed research; LD and JAFDF designed data simulation; GN and VD wrote the code; LD and MQC analyzed data; LD and MQC led the writing. All authors edited and reviewed the text.

